# Flexible tapping synchronization in macaques: dynamic switching of timing strategies within rhythmic sequences

**DOI:** 10.1101/2025.03.27.645824

**Authors:** Ameyaltzin Castillo-Almazán, Oswaldo Pérez, Luis Prado, Nori Jacoby, Hugo Merchant

**Author notes:** Correspondence: Dr. Hugo Merchant.

## Abstract

The ability to synchronize bodily movements with regular auditory rhythm across a broad range of tempos underlies humans’ capacity for playing music and dancing. This capability is prevalent across human cultures but relatively uncommon among non-human species. Recent research indicates that monkeys can predictively synchronize to regular, isochronous metronomes, exhibiting a preference for visual rather than auditory sequences. In this study, we trained macaques to perform a visual synchronization tapping task, testing their synchronization abilities over a wide tempo range and characterizing their precision and accuracy in timing intervals throughout rhythmic sequences. Additionally, we investigated whether the macaques employed priors or error correction strategies to maintain synchrony with the metronome. Our findings demonstrate that, following sufficient training, macaques exhibit a remarkable capability to synchronize across diverse tempos. Through an inference model analysis, we identified two distinct timing control strategies used by the macaques: an initial strong regression-to-the-mean effect transitioning dynamically into a more precise error correction approach at their preferred tempo. These results provide compelling evidence that primates possess sophisticated rhythmic timing mechanisms, effectively leveraging internal and external cues to regulate their tapping behavior according to task demands.

**NEW & NOTEWORTHY:** We trained macaques in a visual synchronization tapping task and using an inference model we found that they are highly precise and accurate in their rhythmic performance. The model also captured a dynamic switch in behavior. At the beginning of the task, the monkeys used their prior knowledge about the interval statistics and then switched to an error correction mechanism to generate intervals depending on the previously produced duration.

## INTRODUCTION

Movement synchronization with external rhythms is a fundamental behavior of modern human life. We dance with a partner at parties or gym classes, march in parades, or play ensemble music with precision and emotional expressivity (Demos and Palmer 2023; Merchant et al. 2015a). These behaviors are highly coordinated, predictive, and demand the multimodal integration of the incoming flow of sensory information (Lenc et al. 2021). Humans show large time flexibility during synchronization perceiving and adapting to musical meter with high dexterity (London 2002). Furthermore, high synchrony is pleasurable and fosters affiliation between partners (Hove and Risen 2009). In music, rhythmic synchronization in an ensemble or orchestra mainly depends on the auditory modality. However, co-performers use visual signals, such as head movements, to provide timing cues (Davidson 2012; Goebl and Palmer 2009), and conductors use dynamic spatiotemporal visual signals to organize the rhythmicity of musicians’ movements (Luck and Sloboda 2009; Pérez et al. 2023). Hence, while humans synchronize more accurately to auditory stimuli rather than flashing rhythmic stimuli, such a metronome, their synchronization prediction and accuracy improve when synchronizing to moving visual stimuli that exhibit sudden bouncing (Gan et al. 2015; Hove et al. 2010, 2013; Iversen et al. 2015; Pérez et al. 2023). It has been suggested that the auditory-visual asymmetry is eliminated when using visually moving metronomes, as the visual motion engages the time-to-collision mechanism underlying interceptive action control or collision avoidance (Li et al. 2022; Merchant et al. 2004a; Merchant and Georgopoulos 2006). Thus, the parieto-premotor system that is recruited for encoding time-to-contact for discrete events, could also be involved in the internal representation of periodic collision points, efficiently driving the motor system for beat-based timing in humans (Balasubramaniam et al. 2021; Iversen et al. 2015; Mendoza and Merchant 2014). In contrast, monkeys are better at synchronizing to visual flashing than auditory metronomes (Betancourt et al. 2023; Merchant et al. 2013a; Zarco et al. 2009) (Betancourt et al. 2023; Merchant et al. 2015a; Zarco et al. 2009). In this case, the hypothesis postulates that the connectivity within the audiomotor system is less developed in monkeys than in humans (Honing and Merchant 2014; Merchant and Honing 2014), despite the highly reciprocal connections in the visual parieto-premotor system in both species (Bufacchi et al. 2023; Caminiti et al. 2010; Merchant et al. 2011). Therefore, the bias towards visual metronomes in macaques is due to their strong visuomotor system, which efficiently extracts rhythmic events from the optic array (Merchant et al. 2001, 2004b, 2023).

During tapping synchronization to a metronome, the subjects need to detect the metronome to produce an internal representation of its tempo, which predicts the rhythmic events that follow, and to generate timing motor commands in anticipation of predicted future beats (Betancourt et al. 2023; Merchant and de Lafuente 2024). Classical tapping studies have identified a preferred rhythmic tempo corresponding to the spontaneously produced duration when subjects rhythmically respond without external cues. This spontaneous or preferred tempo ranges from 500 to 700 ms in human adults (Fraisse 1978; McAuley et al. 2006; Tranchant et al. 2022). Therefore, the internal neural mechanism for rhythmic timing must have a preferred tempo that changes between subjects (Garcia-Saldivar et al. 2024). Error correction in the tapping sequence is another crucial aspect of sensorimotor synchronization, facilitating the compensation of interval production. In error correction, a short interval is typically succeeded by a long interval and vice versa, thereby preventing significant error accumulation with respect to the isochronous stimuli (Jantzen et al. 2018; Repp and Penel 2004). Error correction is usually captured using the negative lag 1 autocorrelation of intervals produced in a large tapping sequence, which determines the alternating serial dependence of long-short or short-long intervals (Hove et al. 2010; Semjen and Ivry 2001). Recently, we computed the correlation of pairs of adjacent produced intervals during a synchronization task to identify the neural correlates for the error correction for different tempos and modalities in monkeys (Betancourt et al. 2023). Finally, the duration of produced intervals during a tapping task can follow the regression towards the mean or the bias effect, characterized by overestimation of shorter intervals and underestimation of longer target intervals, and with an indifference interval that corresponds to the intermediate duration associated with perfect timing accuracy (Jones and Mcauley 2005; Pérez and Merchant 2018). The bias effect is larger when the input information is noisy, and the subjects use their prior knowledge on the statistics of the experimental stimuli. Conversely, this effect is smaller when there is no uncertainty because the measurements of the input stimuli are very robust (Donnet et al. 2014; Pérez and Merchant 2018; Petzschner et al. 2015).

Here we trained macaques in a visual synchronization tapping task to test whether monkeys can synchronize to a wide range of tempos, characterizing the precision and accuracy of the produced intervals and measuring whether they used priors or error correction to keep their taps in synchrony with the metronome. In addition, we developed an inference model which fully explained the key behavioral features of the monkey’s tapping. With this framework of statistical inference, we found that macaques utilized one of two strategies to control rhythmic timing during tap synchronization. At the beginning of the trials, they showed a large regression towards the mean effect, and then dynamically changed to an error correction mechanism once they had timing information about the metronome and the previously produced interval.

## MATERIALS AND METHODS

### Monkeys

Two Rhesus monkeys (*Macaca mulatta*), referred to as M1 (female, 5.5 kg body weight [BW]) and M2 (male, 12.5 kg BW), were used. All animal care, housing, and training procedures were approved under protocol 0.81H from the Bioethics Research Committee of the Instituto de Neurobiología, Universidad Nacional Autónoma de México. The protocol followed the 3Rs and conformed to the principles outlined in the Guide for Care and Use of Laboratory Animals (NIH, publication number 85-23, revised 1985) and the Official Mexican Standard NOM-062-ZOO-1999, Technical Specifications for the Production, Care, and Use of Laboratory Animals. Researchers and animal care professionals monitored the monkeys daily to assess their health and welfare conditions.

### Apparatus

Monkeys were seated comfortably in a primate chair facing an 8 × 8 inch LED matrix (Adafruit NeoPixel, with a refresh rate of 1000 Hz, located 56 cm from their eyes). They placed their right hand on a key with an infrared optical sensor (Balluf BOS 11K, response time ≤ 1 ms) and tapped a metal button equipped with a proximity sensor (NBB2-12GM50-E2-V1). Monkeys performed the tasks in a soundproof and quiet room. Tucker-Davis Technologies (TDT) hardware (RZ2) and software (Pynapse) were used to deliver stimuli and collect behavioral data at 12 kHz.

### Isochronous synchronization tapping task (ISTT)

The two monkeys were trained in this task. A trial started when the monkey held a lever during the presentation of a yellow background and three successive flashing red square that acted as metronome (Figure 1A). The disappearance of the yellow background was the go-signal to tap a button seven times in synchrony with the red-square metronome, generating seven inter-response intervals, called produced intervals (*t*_*p1*_ to *t*_*p7*_). The inter-onset interval (IOI) of the metronome was randomly selected from 16 target durations, *t*_*d*_ (400 - 1150 ms) in trial blocks. Each block consisted of five training trials, where the monkey adapted to the new tempo, and twenty correct testing trials. The monkeys completed between 10 and 12 blocks in a day. The monkeys were rewarded with a few drops of water when two conditions were met: the produced intervals during the trial did not deviate by more than 22% from the instructed interval, and all asynchronies between stimuli and taps were less than 22% of the instructed interval. We allowed 50 ms (25 ms positive and 25 ms negative) of extra time in the asynchronies of the first tap, since the monkey was moving from the key to the bottom. Finally, the reward was doubled or quadrupled when the absolute asynchronies were from 50-100 ms or 0-50 ms, respectively, to promote predictive timing.

**Figure 1.**
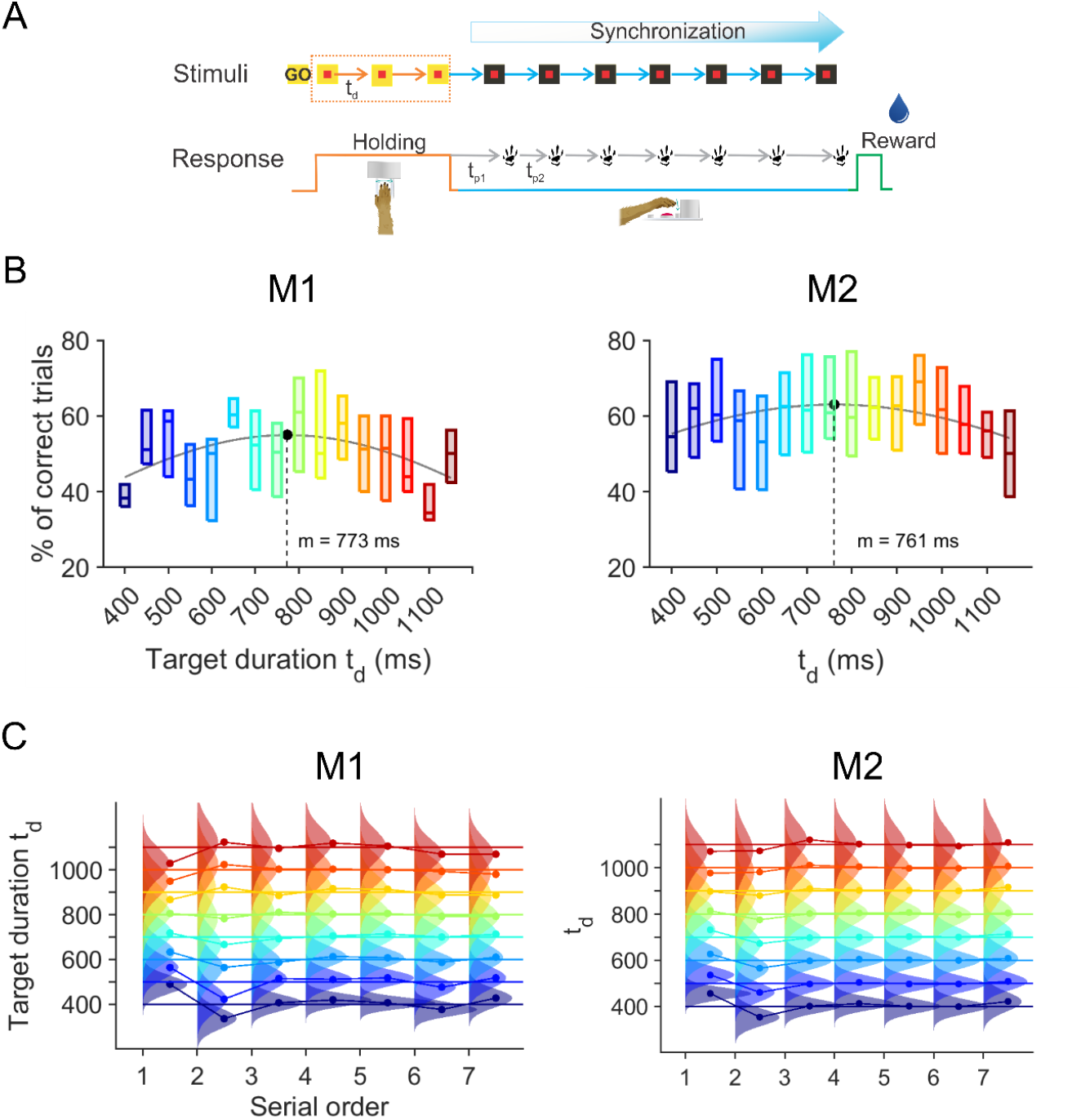
Isochronous tapping synchronization task and distributions of produced intervals (*t*_*p*_). **A**, Experimental setup and trial structure. A trial started when the monkey held a lever during the presentation of a yellow background and three flashing red squares that acted as a metronome. The disappearance of the yellow background was the go-signal to tap a button seven times in synchrony with the red square metronome, producing seven intervals (*t*_*p1*_ to *t*_*p7*_). The inter-onset interval (IOI) of the metronome was randomly selected from 16 target durations, *t*_*d*_ (400 – 1,150 ms). The monkeys completed five training trials and 20 correct test trials of one interval before pseudorandomly switching to a block of trials of a different interval. **B**. Percentage of correct trials as a function of target duration (*t*_*d*_). Each boxplot corresponds to the median (central line) and the 25th (bottom edge) and 75th percentiles (top edge) of a group of blocks for each *t*_*d*_. *t*_*d*_ are color coded. The gray line is the fitting of a Gaussian function with a mean of 773 ms for M1 monkeys and 761 ms for M2 monkeys. **C**, Distributions of the produced intervals *t*_*p*_ as a function of serial order (SO1 - SO7) for eight of the sixteen *t*_*d*_ that were selected every other tempo (from the original sixteen) and are color coded. The mean *t*_*p*_ is depicted as a dot.

### Monkey training

Macaques were trained following operant conditioning techniques (Merchant et al. 2003, 2011; Naselaris et al. 2006a, 2006b). They received normal food rations, were water-deprived, and received most of their water intake during the training and testing sessions. The animals worked 5 days a week, averaging 2 hours per day, and completed around 200 correct trials every day, with a total liquid intake of 150–300 ml during performance. Weight was strictly controlled; supplementary fluids were provided when monkeys did not reach their minimum water volume in a day of performance.

At the beginning of the training, the monkeys received a reward whenever they held the lever for a few seconds. Subsequently, they learned to hold the lever while two flashing red squares were presented and to push the button once the LED background turned black and the third red square was displayed. The target duration for the three stimuli was set at 450 ms. To consider a tap in synchrony with the visual stimulus, we established a large temporal window around the stimulus onset (±60% of IOI). This initial training phase lasted three weeks. Once the monkeys understood that they needed to hold the lever and push the button (more than 60% of the trials for a session over one week), we varied the tempo of the metronome on a trial-by-trial basis ranging from 450 to 850 ms, differing by 50 ms. Critically, the reward was delivered only on trials with asynchronies close to zero (± 150 ms), with extra rewards for smaller asynchronies. The number of stimuli and required taps in a trial only increased when the monkeys were stable at producing precise asynchronies. This training strategy was essential for developing the monkeys’ predictive behavior regarding the timing of the metronome. Once they reached 65% of correct trials in the full version of the isochronous task (seven taps, interval between 450 and 850 ms), we collected the data for this study using the sixteen tempos that covered a wider range of tempos. M1 monkeys performed the task consistently with seven taps after 10 months of training, whereas M2 monkeys achieved this in 5 months.

### Data analysis

All offline data processing and analyses were conducted using MATLAB (2022b, MathWorks). We evaluated the monkeys’ performance during the ISTT by analyzing their tapping response times, acquired at 12 kHz from the TDT and down sampled at 1200 Hz. The series of seven taps produced during each trial defined seven produced intervals (*t*_*p*_) in synchrony with the metronome. The metronome IOI was referred to as target duration (t_d_). For each session, we computed the mean constant error (CE) and temporal variability (TV) of each of the seven *t*_*p*_ across the trials in the same experimental condition. CE is defined as *t*_*p*_ – *t*_*d*_ and represents a measure of timing accuracy. TV is a measure of timing precision and corresponds to the standard deviation of *t*_*p*_.

### Slope analysis

The slope method is a classical timing model that utilizes a linear regression between TV as a function of target duration (*t*_*d*_) to arrive at a generalized form of Weber’s law (see Figure 2A). The linear model follows the equation *TV*(*t*_*d*_) = *intercept*_*TV*_ + *slope*_*TV*_ ∗ (*t*_*d*_ − t_0_). The resulting slope (*slope*_*TV*_) is associated with the time-dependent process, capturing the scalar property of interval timing. The intercept (*intercept*_*TV*_) is related to the time-independent component, which is the fraction of variance that is relatively invariant across interval durations. This component is associated with sensory detection and processing, decision making, memory load, and/or motor execution (Ivry and Hazeltine 1995; Merchant et al. 2008a; Zarco et al. 2009). As a convention, we computed the *intercept*_*TV*_ at the intermediate target interval of *t*_0_ = 775*ms* instead of at 0 *ms*, which represent the indifference interval, namely, the interval where there is no timing error.

**Figure 2.**
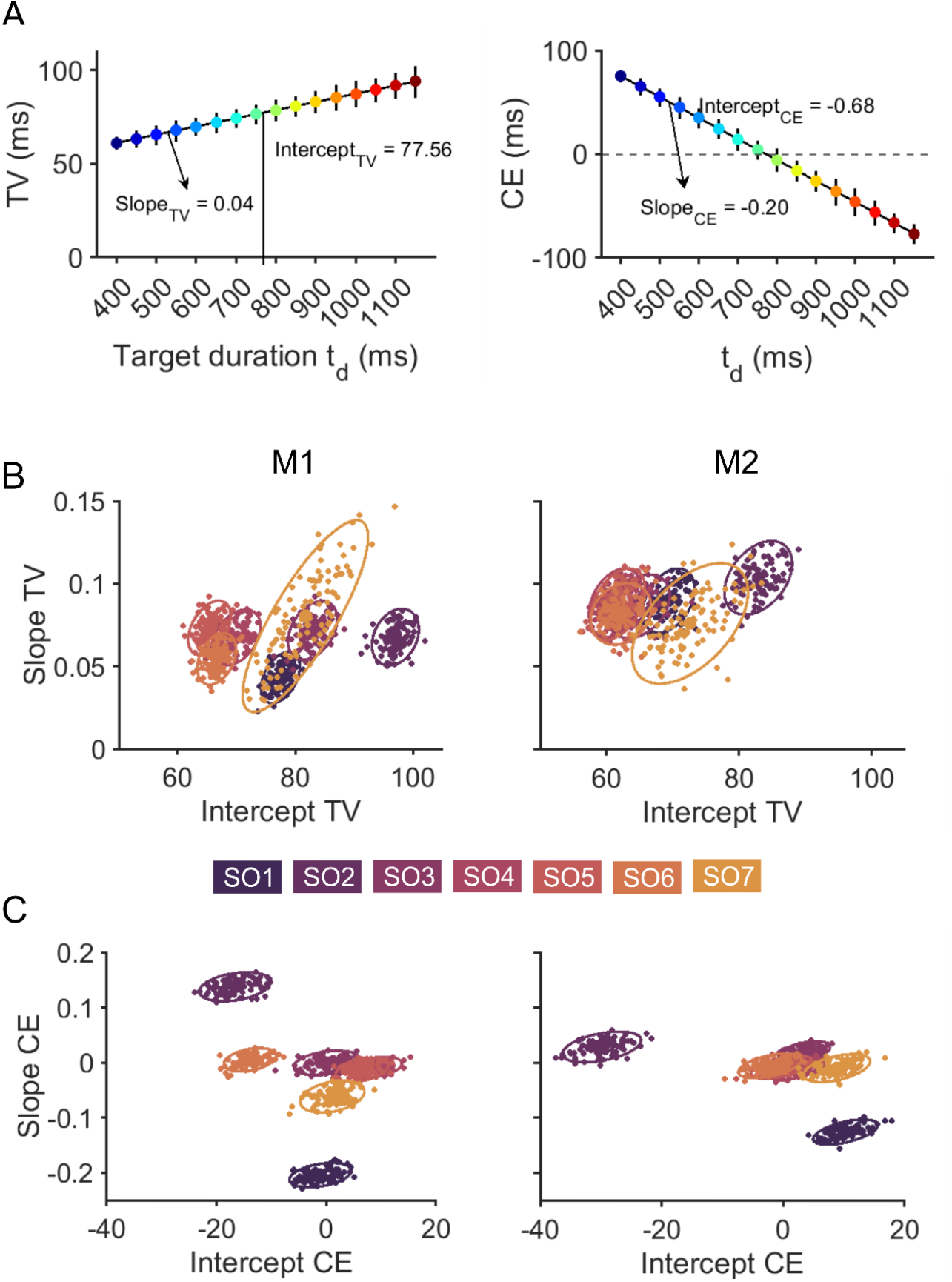
Timing precision and accuracy. **A**, Temporal variability (TV) and constant error (CE) of the first serial order (SO1) of M1 monkeys as a function of *t*_*d*_. We fitted a linear regression for both parameters and determined the corresponding intercept (Intercept_TV_ = 77.56 ms; Intercept_CE_ = -0.68ms), and slope (Slope_TV_ = 0.04 ms; Slope_CE_ = -0.20 ms). **B**, Slope_TV_ is plotted as a function of Intercept_TV_ for each serial order for both monkeys. Serial order is color-coded, from dark violet (SO1) to yellow (SO7). **C**, Slope_CE_ is plotted as a function of intercept_CE_ for each serial order for both monkeys. Same color code as in B.

### Inference model

To investigate the timing processing across serial orders in the synchronization task, we employed a modified version of the Bayesian Model proposed by Perez et al., 2023. This comprehensive model encompasses four key processing stages: 1) the internal measurement *t*_*m*_ of the input time *t*_*s*_, 2) the internally estimated time *t*_*e*_, 3) the produced time *t*_*p*_, and 4) the feedback processes within the rhythmic sequence. By incorporating these interconnected stages, the model is able to capture the essential properties of the bias and scalar effects observed in timing synchronization paradigms. Specifically, we model the likelihood probability *p*(*t*_*p*_|*t*_*d*_, t_*p*−1_), that reflects the probability of observing the produced interval *t*_*p*_, given the target interval *t*_*d*_ and the previously produced interval *t*_*p*−1_. This probability is calculated considering the internal processes of measurement, estimation, production, and feedback, following marginal probability:

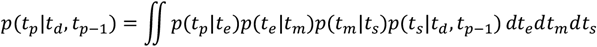

For the internal measurement process, we assume that the conditional probability *p*(*t*_*m*_|*t*_*s*_) follows a normal distribution with a mean *t*_*s*_ and standard deviation *σ*_*m*_, i.e., *p*(*t*_*m*_|*t*_*s*_)∼*N*(*t*_*s*_, *σ*_*m*_). Note that *σ*_*m*_ is a parameter in the model that represents the amount of noise added to the internal measurement.

In the estimation stage, we assume that the observer computes a time estimate *t*_*e*_ that is linearly dependent on *t*_*m*_, using the function *t*_*e*_ = *f*(*t*_*m*_) = *b*_*e*_ + *m*_*e*_(*t*_*m*_ − 775). 775 ms is the intermediate interval of the sample durations. Both *b*_*e*_ and *m*_*e*_ are model parameters and represent the bias and slope of the relationship between the measured *t*_*m*_ and the estimated time *t*_*e*_, respectively. This model is a simplification of the maximum likelihood estimation of Perez et al., 2023. The relationship between *t*_*m*_ and *t*_*e*_ is deterministic, so *p*(*t*_*e*_|*t*_*m*_) = δ(*t*_*e*_ − *f*(*t*_*m*_)).

In the production stage, we assume that the conditional probability *p*(*t*_*p*_|*t*_*e*_) follows a normal distribution with mean *t*_*e*_ and standard deviation *w*_*p*_*t*_*e*_, i.e., *p*(*t*_*p*_|*t*_*e*_)∼*N*(*t*_*e*_, *w*_*p*_*t*_*e*_). *w*_*p*_ is another model parameter, and the linear relationship *w*_*p*_*t*_*e*_ allows us to mimic the scalar property of the produced intervals in synchronization tasks.

Finally, the feedback process estimates the input time by comparing the target time *t*_*e*_ and the previously produced time *t*_*p*−1_ following the equation *t*_*s*_ = *g*(*t*_*d*_, t_*p*−1_) = t_*d*_ + *λ*(*t*_*p*−1_ − t_*d*_). *λ* is the final parameter of the model and represents the weight of the feedback. *λ* takes values between -1 and 1, where *λ* = 1 implies the reproduction of the previous interval, i.e., *t*_*s*_ = t_*p*−1_; *λ* = 0 means no feedback, i.e., *t*_*s*_ = t_*d*_; while *λ* = −1 represents error correction. Therefore, *p*(*t*_*s*_|*t*_*d*_, t_*p*−1_) = δ(*t*_*s*_ − *g*(*t*_*d*_, t_*p*−1_)).

By integrating the marginal probability with respect to *t*_*e*_ and *t*_*s*_ and considering the delta probabilities *p*(*t*_*e*_|*t*_*m*_) and *p*(*t*_*s*_|*t*_*d*_, t_*p*−1_), the posterior probability can be simplified as:

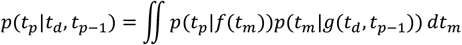

To end, we used *p*(*t*_*p*_|*t*_*d*_, t_*p*−1_) to maximize the likelihood probability for *σ*_*m*_, *b*_*e*_, *m*_*e*_, *w*_*p*_ and *λ* across all pairs of sequentially produced intervals. The maximization was carried out using a gradient-based method in MATLAB. For each of the sixteen *t*_*d*_ we collected 400 and 200 correct trials in M1 and M2 monkeys, respectively. From these totals we randomly selected 50 trials, on which we performed the slope analysis and the model fitting, resulting in 100 iterations of analysis.

### Statistical analysis

For each monkey we performed a repeated measures analysis of variance (rmANOVA) using CE or TV as independent parameters and the target duration (*t*_*d*_: 400, 1150, 16 levels) and serial order of *t*_*p*_ (7 levels) as “within subjects” factors. In addition, we carried out rmANOVA models using the coefficients from the slope analysis (slope_TV_, intercept_TV_, slope_CE_, intercept_CE_) and Bayesian model (*σ*_*m*_, *b*_*e*_, *m*_*e*_, *w*_*p*_ and *λ*) as the independent factor and the serial order of *t*_*p*_ (7 levels) as the “within subjects” factor.

Greenhouse-Geisser corrections were applied to the ANOVA factors’ p-values with a statistical significance cut-off set at p < 0.05. Finally, pairwise comparisons between levels of the ANOVA factors were evaluated by means of two-tailed paired t-tests, with Bonferroni correction applied (cut-off of p < 0.05).

## RESULTS

We trained two monkeys in an ISTT (see Methods). The animals sat in a primate chair and started a trial by holding a lever while a yellow background with flashing red squares, acting as a metronome, was displayed on a LED matrix (Figure 1A). The disappearance of the yellow background was the go-signal to tap a button seven times in synchrony with the red-square metronome, generating seven inter-response or produced intervals (*t*_*p*_). The animals received a liquid reward if their accuracy and asynchronies fell below a specified error threshold (see Methods). Sixteen IOIs (400 – 1,150 ms), referred to as target durations (*t*_*d*_), were presented pseudorandomly in blocks with five training trials during which the monkey adapted to the new tempo and with then twenty correct testing trials.

Monkeys were trained to predictively tap into a sequence of isochronous stimuli, with the number of elements increasing progressively once the animals reached 65% of correct trials for the initial range of tempos (450-850 ms; see Methods). Data for this paper was collected when monkeys were able to synchronize accurately and precisely within a window of 22% of the *t*_*d*_. M1 monkeys performed the task consistently with seven taps after 10 months of training, while M2 monkeys took 5 months. Nevertheless, the percentage of correct trials varied across *t*_*d*_. Both animals showed a higher percentage of correct trials at intermediate tempos, achieving best performance at 773 and 761 ms for M1 and M2 monkeys, respectively (Figure 1B). This interval is probably an expression of the internal preferent tempo of the animals and is not due to the training *t*_*d*_, since in the training the interval ranged from 450 to 850 ms. Thus, produced intervals that depend on the training process would likely to be within that range and not around 770 ms as observed. A one-way ANOVA showed significant main effects in *t*_*d*_ for both animals, supporting the notion of a preferred tempo at which monkeys performed the task more consistently (M1: F(15, 167) = 2.6, *p*<0.0016; M2: F(15,498) = 2.13, p<0.0077).

Figure 1C shows the distributions of correct *t*_*p*_ for eight of the 16 *t*_*d*_ (every other tempo) as a function of the serial order in the synchronization task. There is an increase in the width of the distributions across target durations. In addition, it is evident that the first produced interval in the sequence showed a significant regression towards the mean, with overestimation for short intervals and underestimation for long intervals. The second produced interval showed the opposite effect, and the last five *t*_*p*_ showed a duration distribution that was near the target duration. To parametrically characterize this behavior, we used three performance measures: temporal variance (TV), defined as the standard deviation of *t*_*p*_ (Figure 2A); constant error (CE), which is the difference between produced *t*_*p*_ and target *t*_*d*_ (Figure 2B); and the correlation of the consecutive produced intervals in the seven-interval sequence (Figure 3). The first metric assesses timing precision, the second evaluates timing accuracy, and the third is an index for error correction in synchronization.

**Figure 3.**
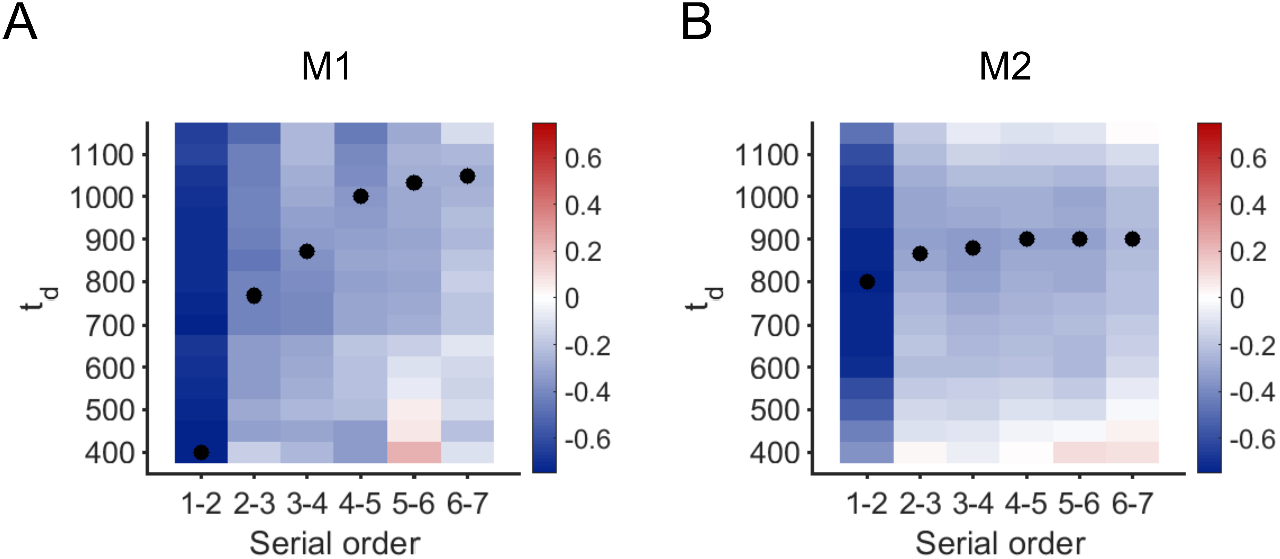
Heatmaps of the correlation of the *t*_*p*_ of consecutively produced intervals (e.g., 1-2 or 2-3) for the 16 *t*_*d*_ for M1 in A and M2 in B. Negative correlations are blue, positive correlations are red. The scale on the heatmap is on the right. Black dots represent the most negative correlation (all p < 0.0001) for each pair of consecutive intervals and for each *t*_*d*_.

### Tapping precision

The TV increased linearly as a function of *t*_*d*_, following the scalar property of timing (Merchant et al. 2008a; Pérez and Merchant 2018, Figure 2A). An rmANOVA showed significant main effects for *t*_*d*_ and serial order, as well as for the *t*_*d*_ x serial order interaction (see Table 1). We performed a linear regression between TV as a function of *t*_*d*_ to arrive at a generalized form of Weber’s law (see Figure 2A). The resulting slope (slopeTV) is associated with the time-dependent process, capturing the scalar property of interval timing. The intercept (interceptTV) is related to the time-independent component, which is the fraction of variance that remains relatively constant across interval durations. It is associated with sensory detection and processing, memory load, and/or motor execution (Ivry and Hazeltine 1995; Merchant et al. 2008a). We computed the interceptTV at the intermediate target interval of 775 ms instead of at 0 ms, as typically done in linear regression (Figure 2A). These analyses were conducted in batches of 50 randomly selected trials for a total of 100 iterations (dots with ellipses in Figure 2C,D). The time-independent component was stable across serial order elements (Figure 2B), with values between 0.05 and 0.1 in the two animals. However, the corresponding rmANOVA showed a significant effect for serial order (Table 1). In addition, InterceptTV showed significantly larger values in the second produced interval and a wider range in the last interval of the sequence, along with substantial statistical changes (Table 1).

**Table 1.**
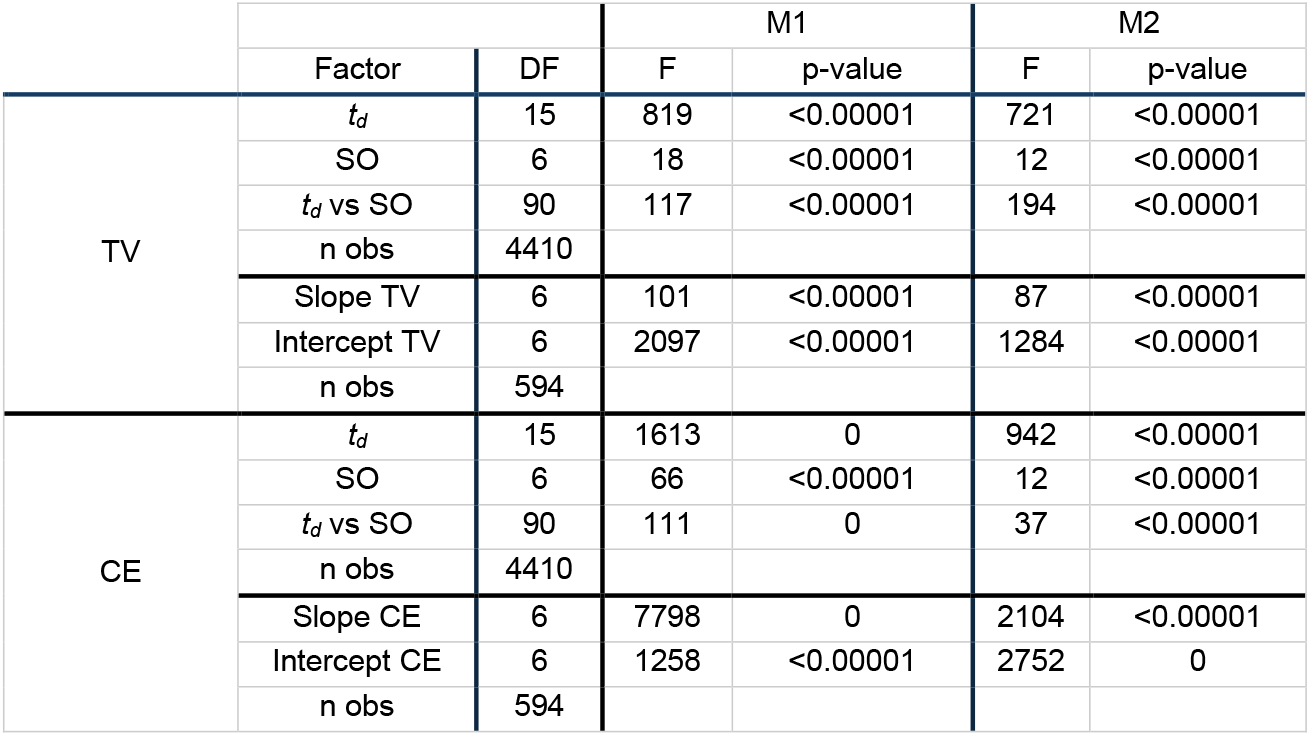
Results of repeated measures ANOVA (rmANOVA) for TV (temporal variability) and CE (constant error).

|  |  |  | M1 |  | M2 |  |
| --- | --- | --- | --- | --- | --- | --- |
|  | Factor | DF | F | p-value | F | p-value |
| TV | $t_d$ | 15 | 819 | <0.00001 | 721 | <0.00001 |
|  | SO | 6 | 18 | <0.00001 | 12 | <0.00001 |
| | $t_d$ vs SO | 90 | 117 | <0.00001 | 194 | <0.00001 |
|  | n obs | 4410 |  |  |  |  |
|  | Slope TV | 6 | 101 | <0.00001 | 87 | <0.00001 |
|  | Intercept TV | 6 | 2097 | <0.00001 | 1284 | <0.00001 |
|  | n obs | 594 |  |  |  |  |
| CE | $t_d$ | 15 | 1613 | 0 | 942 | <0.00001 |
|  | SO | 6 | 66 | <0.00001 | 12 | <0.00001 |
| | $t_d$ vs SO | 90 | 111 | 0 | 37 | <0.00001 |
|  | n obs | 4410 |  |  |  |  |
|  | Slope CE | 6 | 7798 | 0 | 2104 | <0.00001 |
|  | Intercept CE | 6 | 1258 | <0.00001 | 2752 | 0 |
|  | n obs | 594 |  |  |  |  |

### Tapping accuracy

Using CE as a dependent variable, the rmANOVA showed significant main effects for *t*_*d*_ and serial order, as well as in the *t*_*d*_ x serial order interaction (Table 1). Figure 2B shows CE as a function of *t*_*d*_ for the first produced interval with the stereotypical bias effect of overestimation for short intervals and underestimation for long intervals. We defined two measures to quantify this effect using linear regression: the intercept at 785 ms (interceptCE) and the regression slope (slopeCE; see Figure 2E). The former is proportional to the indifference interval, namely, the interval where there is no error in timing (Gámez et al. 2018; Jones and Mcauley 2005). The latter corresponds to the magnitude of the bias effect (i.e., the larger the negative slope, the larger the regression towards the mean) (Pérez and Merchant 2018). We plotted the slopeCE as a function of the interceptCE for each produced interval in the task sequence. According to the tapping distribution illustrated in Figure 1B, there is a large regression towards the mean (with a large negative slope CE) in the first produced interval, followed by a positive slopeCE and a negative interceptCE in the second produced interval. In contrast, both slopeCE and interceptCE are stable and close to zero for the rest of the serial order elements. The rmANOVA showed a significant serial order effect in both parameters and both monkeys (Table 1).

### Control strategy for the production of rhythmic sequences

The correlation showed a large negative value between the initial and the second produced *t*_*p*_ across all *t*_*d*_ (Figure 3). It also showed a strong tendency to be close to zero from the 2^nd^ to the 7^th^ produced intervals for the shorter and longer *t*_*d*_, and displayed negative values for the intermediate *t*_*d*_ across same serial order elements. In fact, for each pair of consecutive intervals, the *t*_*d*_ with the most negative correlation is represented as a black dot, and for M2 it converges at around 850 ms (Figure 3B).

These effects are less prominent but still present in M1 (Figure 3A).

### An inference model for tapping synchronization

We adapted the four-stage Bayesian model by Perez et al. (2023) to our synchronization task (Figure 4; see Methods). Indeed, we modeled the observed produced intervals *t*_*p*_ and target interval target *t*_*d*_ using the likelihood probability (p), which reflects the probability of observing the *t*_*p*_ given the target interval *t*_*d*_. Our model comprised four stages: measurement, estimation, production, and feedback. In the initial measurement stage, the physical interval (*t*_*s*_) is transformed by a noisy observer into measured time (*t*_*m*_) using a Gaussian distribution centered on *t*_*s*_ and a standard deviation *σ*_*m*_. Then, the observer computes the maximum likelihood estimator (*t*_*e*_) as the maximum of the posterior probability, which is proportional to the product of the likelihood function of the measured time *p*(*t*_*m*_|*t*_*s*_) and bias property function. The latter is a linear function with a slope *m*_*e*_ and the constant *b*_*e*_ that represents the internal magnitude of the bias effect.

**Figure 4.**
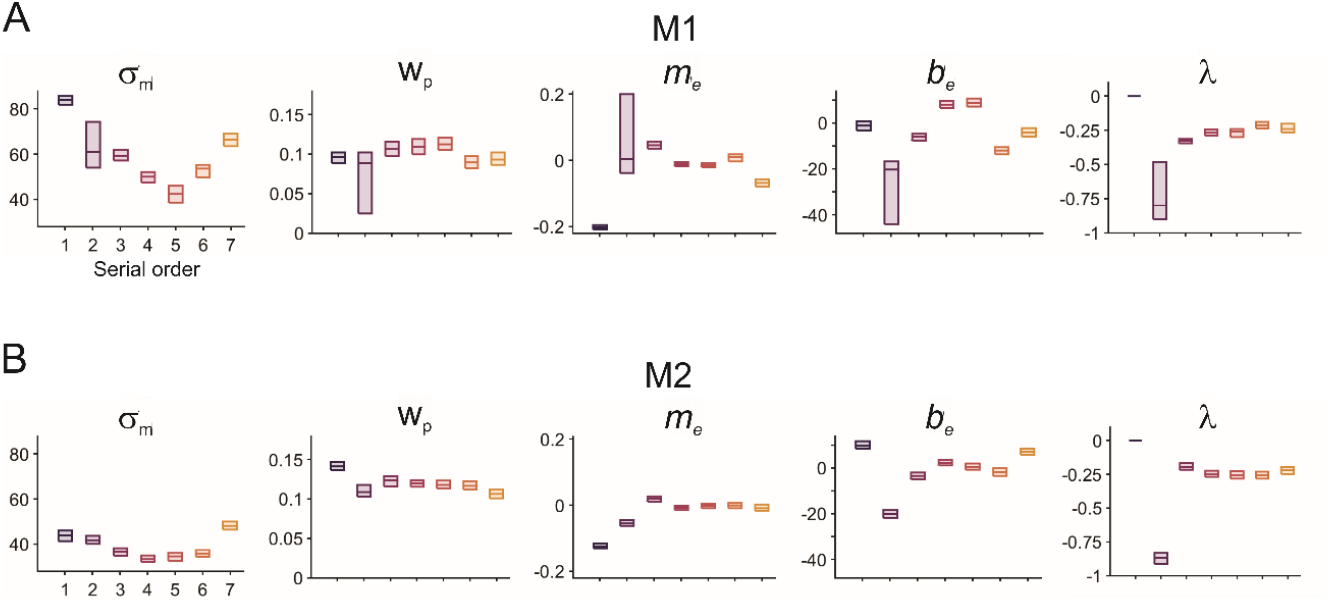
Parameters of the inference model across the serial orders for M1 in A and M2 in B. *σ*_*m*_ corresponds to the time measurement noise, *m*_*e*_ and *b*_*e*_ are the slope and the constant of the bias effect linear regression, *w*_*p*_ corresponds to the scalar property in the produce interval, and *λ* is the feedback coefficient.

The third stage uses *t*_*e*_ to compute the produced interval *t*_*p*_ utilizing the conditional dependence of *t*_*p*_ on *t*_*e*_ as *p*(*t*_*p*_|*t*_*e*_), which adds noise to the production phase through a Gaussian distribution, whose standard deviation is *w*_*p*_*t*_*e*_. Hence, *w*_*p*_*t*_*e*_ increases for longer estimated intervals, simulating the scalar property of timing. The fourth stage of the model captures the structure of the intervals produced sequentially during synchronization. Thus, we calculated the difference between the previously produced interval (*t*_*p*−1_) and the physical interval (*t*_*d*_) and determined how this difference influences the input time as *t*_*s*_ = t_*d*_ + *λ*(*t*_*p*−1_ − t_*d*_) with a *λ* weight. *λ* = 0 in the first produced interval. In the following intervals, *λ* depends on how much error correction adjusts the actual *t*_*p*_, with values below zero for strong error correction since there is a negative feedback loop (Figure 4C, *λ* top). In contrast, *λ* tends to be positive when consecutive intervals are similar, producing a positive loop in the sequence (Figure 4C, *λ* bottom). Please see Figure 4c of Perez et al., 2023 to understand the link between the model parameter and the scalar and bias properties.

For each monkey, we fitted the model parameters, namely, *σ*_*m*_, *m*_*e*_, *b*_*e*_, *w*_*p*_, and *λ* across serial order elements of the task for each of the 100 iterations of 50 randomly chosen trials. Goodness-of-fit values were large, with significant correlations between the predicted and actual data for both monkeys (Pearson correlations with a p < 0.0001). Figure 4 shows the coefficients of the model across the serial orders for the two animals. rmANOVAs showed significant effects of serial order on the five parameters of the model with similar sequential effects in both monkeys (Table 2). The time measurement noise (*σ*_*m*_) showed large values for the first and last produced interval in the sequence, suggesting larger processing noise when starting entrainment and expecting a reward at the end of the trial. The slope of the bias effect (*m*_*e*_) was negative for the first produced interval and approached zero in the rest of the intervals, indicating a strong regression towards the mean only at the beginning of the trial. *b*_*e*_ was close across serial orders with the exception of the second produced interval, which exhibited negative values, demonstrating that monkeys showed an indifference interval close to 775 ms. The *w*_*p*_ was similar across serial orders, indicating a stable scalar properly. Finally, *λ* was negative for all serial orders, although it was close to -1 for the second serial order. This last result supports the existence of a large error correction mechanism during the synchronization, especially in the second produced interval.

**Table 2.**
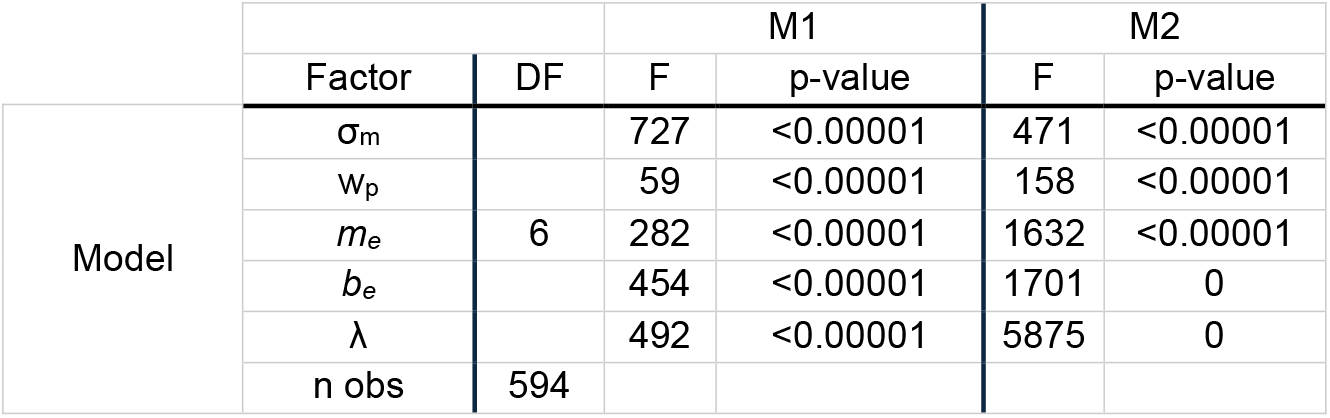
rmANOVA results for the parameters of the inference model.

## DISCUSSION

The results of the present study support three main conclusions. First, macaques show a large flexibility in synchronizing their tapping to visual metronomes over a wide range of tempos. Second, the monkeys started a trial using a strong initial regression towards the mean strategy, then switched to a large error correction mechanism, where the second produced interval had the tendency to have an inverse CE from the first, and finally showed a stable tapping synchronization using a moderate error correction mechanism across the rest of the produced intervals. Lastly, a simple inference model captures all these complex features and can be used as a framework to study the neural basis of rhythm entrainment.

The notion that only humans can perceive and entrain to rhythms has evolved dramatically over the last decade. The original gradual audiomotor hypothesis posits that beat-based timing emerged gradually in primates, peaking in humans due to a sophisticated audiomotor circuit. Macaques exhibit basic features of this timing because of their sophisticated auditory cortex, basal ganglia, and medial premotor system (Merchant and Honing 2014). Since then, studies have shown that primates have complex rhythmic timing. On the one hand, EEG studies in Rhesus monkeys have shown that macaques produce evoked potentials linked to the detection of isochronous auditory patterns (Ayala et al. 2017; Honing 2018; Honing et al. 2012), as well as to subjectively accented 1:2 and 1:3 rhythms from auditory metronomes (Criscuolo et al. 2023). On the other hand, biomusicologists have demonstrated that great apes show rhythmic drumming behaviors (Ravignani et al. 2013) and can learn to execute human-like sequences (Fuhrmann et al. 2014), while lemurs produce in the wild coordinated songs with isochronous and 1:2 rhythmic categories (De Gregorio et al. 2021). Finally, monkeys trained in tapping tasks can flexibly and predictively produce periodic intervals in synchrony with auditory and visual metronomes (Betancourt et al. 2023; Gámez et al. 2018). They can continue tapping without sensory cues (Donnet et al. 2014; Zarco et al. 2009) and can even consistently tap to the subjective beat of music excerpts (Rajendran et al. 2024). Hence, these studies suggest that non-human primates can perceive and synchronize to simple rhythms such as isochrony. A systematic observation in these experiments is that monkeys have a bias towards visual rhythmic stimuli (Betancourt et al. 2023; Gámez et al. 2018), which is the opposite to the human bias towards auditory stimuli (Iversen et al. 2015; Merchant et al. 2008b; Repp 2005). In the present study, we used visual metronomes with a wide range of tempos. The monkeys displayed an astonishing flexibility in synchronizing their tapping with high precision and accuracy. This ability is similar to the human ability to synchronize auditory metronomes with IOIs spanning from 300 to 1200 ms (Repp 2005). To achieve this performance, it is key to reward the tapping behavior of the monkeys with proper phase and tempo matching (Gámez et al. 2018; Rajendran et al. 2024). Hence, these observations suggest that macaques have the brain sensorimotor mechanisms to: (1) extract a rhythm from a continuous stream of sensory events, (2) generate an internal rhythmic signal that predicts the future beat events, and (3) produce motor commands such that movements coincide or slightly anticipate the next rhythm (Betancourt et al. 2023; Mendoza and Merchant 2014). In addition, it is evident that metronome synchronization is not as natural to monkeys as it is to humans, and external rewards with clear performance rules are needed for appropriate rhythmic entrainment (García-Garibay et al. 2016; Merchant et al. 2024; Rajendran et al. 2024; Takeya et al. 2017).

Here we found that the monkeys exhibited a sophisticated organization of sequential tapping across tempos and serial orders. During the first produced interval in the sequence, which corresponds to the time between the go signal and the first button tap after releasing the key, the animals used a large regression towards the mean strategy. The CE as a function of *t*_*d*_ showed a steep negative slope, with significant overestimation for short target intervals and substantial underestimation for long target intervals, as well as an indifference interval close to 775 ms, which corresponds to the midpoint of the tested tempos. Therefore, at the beginning of the trial, the monkeys used a robust prior systematically biased towards the center of the tested distribution of *t*_*d*_, similar to what has been documented in single interval reproduction tasks in humans and monkeys (Jazayeri and Shadlen 2010; Wang et al. 2018). In all these tasks the subjects are cued with a set interval, which they reproduced with a movement (Merchant et al. 2013b; Remington et al. 2018). In contrast, during the next produced interval, the monkeys used a large error correction strategy to synchronize with the metronome. This indicates the strongest negative correlation between consecutively produced intervals with a switch in the sign of the CE between the first and second intervals across *t*_*d*_, showing good correcting behavior, especially in M1 monkeys. As far as we know, this is the first evidence of an immediate shift from a robust regression towards the mean strategy to an error correction mechanism in rhythmic timing. Classical studies on sensorimotor synchronization have focused on the stable phase of tapping, dropping the first produced interval from the analysis and explaining why this phenomenon has not been reported. Nevertheless, recently we found that human subjects switch from error correction during tapping synchronization to a regression towards the mean strategy during the internally driven tapping continuation epoch, without a metronome (Pérez et al. 2023). From the third to the seventh elements of the sequence, the animals used a moderate error correction mechanism with negative correlations between consecutively produced intervals. Notably, the target durations with the most negative correlations between adjacent *t*_*p*_ were close to 900 ms in M1 monkeys and 850 ms in M2 monkeys, supporting the notion that animals have a preferred tempo at which they can correct their rhythmic tapping more robustly. In addition, this tempo is close to the InterceptCE where the animals demonstrated high accuracy in their rhythmic performance. Hence, these results suggest a strong relationship between proper error correction and high rhythmic timing accuracy at a preferred tempo, an association previously observed in humans (Tranchant et al. 2022).

Our inference model could account for the stable scalar property across the rhythmic sequence and detect the optimal change in strategy, starting with the regression towards the mean and switching to an error correction mechanism. This is consistent with previous Bayesian model on timing behavior (Cannon 2021; Jazayeri and Shadlen 2010; Sadakata et al. 2006; Sohn and Jazayeri 2021). Hence, the statistical inference framework can be useful to study the neurophysiological bases of rhythmic timing in monkeys. Recent neurophysiological studies in monkeys indicate that the internal pulse representation during rhythmic tapping to visual and auditory metronomes depends on the neural population dynamics in the medial premotor cortex (MPC). A key property of MPC neurons is their relative representation of beat timing. Cells that encode elapsed or remaining time for a tap show up-down ramping profiles that span the produced interval, with speed scaling as a function of beat (Merchant et al. 2011; Merchant and Averbeck 2017). Furthermore, these cells are recruited in rapid succession, producing a progressive neural activation pattern that flexibly fills the beat duration depending on the tapping tempo, thereby providing a relative representation of the evolution of an interval (Crowe et al. 2014; Mello et al. 2015; Merchant et al. 2015b). The neural cyclic evolution and resetting become more evident when the time-varying activity of MPC neurons is projected into a low-dimensional state space (Gámez et al. 2019; Merchant and Bartolo 2018). The population neural trajectories show the following properties. First, their circular dynamics form a regenerating loop for every produced interval. Second, they converge in a similar state space at tapping times, resetting the beat-based clock at this point. This internal representation of rhythm could be transmitted as a phasic top-down predictive signal to sensory areas (Merchant et al. 2023). Third, the periodic trajectories increase in amplitude and decrease in speed as a function of the length of the isochronous beat encoding the tempo of the metronome (Betancourt et al. 2023). Finally, the scalar property, a hallmark of timing behavior, was accounted for by the variability of the curvilinear radii in neural trajectories (Betancourt et al. 2023; Gámez et al. 2019).

Hence, the neural population trajectories in MPC encode beat, tempo, and variability of rhythmic timing, which are behavioral features captured by our model. In addition, neural trajectories have been linked to error correction during tapping synchronization (Betancourt et al. 2023) and to the timing prior that contributes to the regression towards the mean effect in single interval tasks (Balasubramaniam et al. 2021; Sohn et al. 2019).

In sum, this study provides evidence of a flexible tempo and dynamic rhythmic mechanism in monkeys during tapping synchronization. It suggests avenues for future experiments to explore the neural underpinnings of the dynamic switch from using prior information about task tempos for initial timing to a robust error correction mechanism that is more efficient at the preferred tempo, once the monkeys perceive the metronome and their produced intervals.

## ACKNOWLEDGMENTS

We thank Yaneri Ayala for her fruitful comments on the manuscript and to Jessica Gonzalez Norris for proofreading the paper. We also thank Maria Antonieta Carbajo, Juan Ortíz, and Raul Paulín for their valuable technical assistance.

## AUTHOR CONTRIBUTIONS

H.M. and A.C-A. conceived the experiments. A.C-A. and L.P. collected the psychophysics data. O.P., H.M. and A.C-A. analyzed the data. N.J contributed to the writing of the manuscript providing expert advice. H.M. supervised the project.

